# Three-spined sticklebacks recognise familiar individuals by facial recognition

**DOI:** 10.1101/2024.10.23.619764

**Authors:** Shumpei Sogawa, Izumi Inoue, Satoshi Awata, Koki Ikeya, Kento Kawasaka, Masanori Kohda

## Abstract

Social vertebrates often recognise familiar individuals by facial recognition, a basal cognitive ability by which animals establish stable sociality, including territoriality. The three-spined stickleback (*Gasterosteus aculeatus*), a model species for behavioural studies, is territorial; its ability to visually recognise familiar individuals remains unclear. Here, we report that this species has individual-specific facial colour morph features and can recognise familiar individuals by facial recognition. Territorial neighbours of the same sex established a “dear enemy relationship” with each other. These focal fish were exposed to composite photographic models of four combinations of faces and bodies of familiar neighbours and unknown strangers of the same sex. Focal fish of both sexes frequently attacked photographs of strangers (stranger-face/stranger-body) more frequently than familiar neighbours (neighbour-face/neighbour-body). Furthermore, they attacked composite photographs of stranger-face/neighbour-body more frequently (similar to the stranger model) but less frequently attacked photographs of neighbour-face/stranger-body (similar to the neighbour model). These results suggest that the three-spined stickleback distinguishes familiar neighbours from unknown fish exclusively via facial recognition. The aggressiveness of males was independent of the presence of red nuptial colour on photograph-models. Our findings suggest that this fish controls its aggressiveness against opponent conspecifics in the context of social relationships independent of the sign-stimulus.

## 1. Introduction

The three-spined stickleback (*Gasterosteus aculeatus*), first used for studies of aggressive behaviours by Tinbergen in the 1930s [1, 2, 3, 4], is an important model species in animal behavioural studies (e.g., [5, 6, 7]). During the breeding season, males establish breeding territories and attack intruding conspecific males. Tinbergen interpreted the observations of a series of the classic experiments as an indication that the aggressive behaviour of the three-spined stickleback is instinctively triggered by the male’s nuptial red colour on the belly, which functions as a sign-stimulus [4]. This interpretation has been widely accepted; it is included in current textbooks of general biology and animal psychology as a typical example of a fixed action pattern evoked by sign-stimulus based on “hard-wired” innate releasing mechanisms (e.g., [8, 9]). However, several groups have suggested that the concept of sign-stimulus is oversimplified because the red nuptial colour does not always trigger aggression and aggression does not always require the red sign-stimulus (e.g., [10, 11, 12]).

Indeed, the reciprocal altruism that is difficult to be explained innate releasing mechanisms during predator inspection visits has been documented in the three-spined stickleback [13, 14]. To maintain this sophisticated social relationship with a tit-for-tat strategy, this fish will have a variety of cognitive ability, such as familiar recognition (a type of class revel recognition) and strong memory. These cognitive ability (reciprocal altruism) and innate releasing mechanism will be mutually exclusive to explain the same behaviours. Recently, it is reported that an associative learning study yielded no evidence supporting familiar recognition ability in the three-spined stickleback [15]. The authors claim that this altruistic behaviour will be explained by the concept of shame/border individuality not by individual recognition in the context of Morgan’s Canon. Nevertheless, it is possible that the three-spined sticklebacks in their study did not have no ability of familiar recognition but might simply fail in associative learning. Thus, to consider which process this model animal employs will be quite important. Here we examine their familiar recognition ability from a different angle in an experiment that took into account their ecological characteristic of territorial defence.

Many species of vertebrates from various taxa, including fishes, have breeding territories (e.g., [16]), which are often adjacent to and surrounded by the territories of several conspecific neighbours. In mammals and birds, it has been reported that when a territory boundary has been established between neighbours, it is rarely crossed and they become tolerant of each other (e.g., [17, 18]). This type of social relationship, called a “dear enemy relationship” [19, 20], has been documented in territorial cichlids and guppies [21–27].

It has been suggested that such dear enemy relationships between neighbours occur in the territorial three-spined stickleback [28]. However, their territorial interactions, including this relationship, have not been studied in detail. Analysis of dear enemy relationships provides an opportunity for the study of familiar recognition [23–26]. Cichlid fish that show dear enemy relationships with territorial neighbours visually discriminate familiar neighbours from strangers and can even distinguish between familiar neighbours (i.e., individual recognition) by recognition of individual-specific facial features [23–27, 29, 30]. The three-spined stickleback also has individual-specific facial colour morph characters (figure 1). Therefore, if the territorial three-spined stickleback has dear enemy relationships and distinguish familiar neighbour from stranger depending on their individual-specific faces colour morphs, this fish species may have individual recognition ability similar to cichlids.

**Figure 1.**
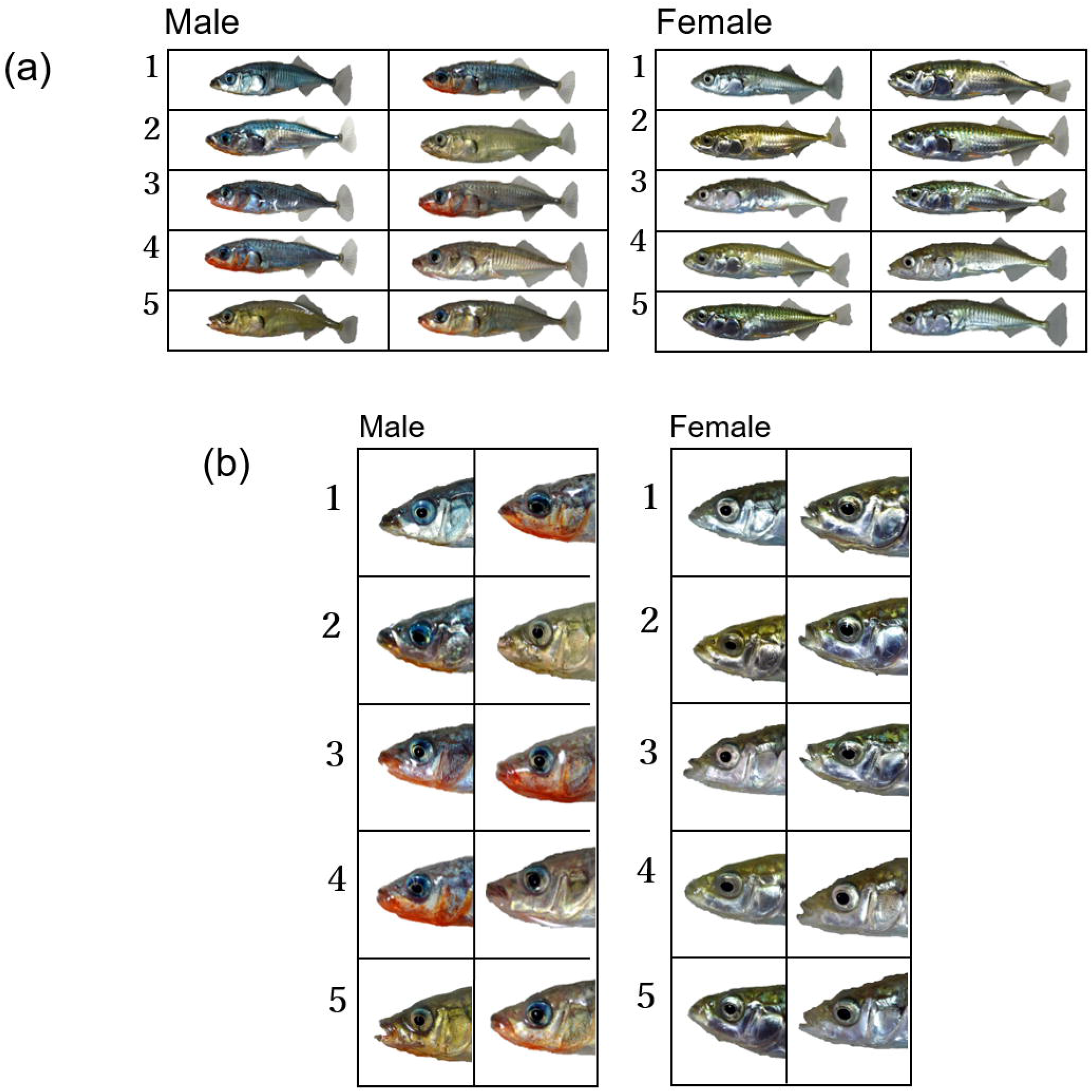
Photographs of 10 males and 10 females of the three-spine sticklebacks (*a*) and their faces (*b*) used for the experiments. Pairs of 1-5 are fish used in making dear enemy relationships in both sexes.

In the present study, we quantitatively examined whether territorial three-spined stickleback can establish dear enemy relationships with their conspecific neighbours, including females that are also potentially territorial under some conditions [29], and whether this fish species can recognise and distinguish familiar neighbours from strangers, especially based on individual-specific facial features. Familiar recognition is one of cognitive abilities essential for maintaining stable social relationships, including territoriality, dominance order, and stable sexual pairs [31]. The process of familiar recognition may include a complex cognitive ability involving recognition, memory, and an inner template or mental image of known individuals [31, 32]. The mental processes of these cognitive abilities generally will be different from the simple innate releasing mechanism [3, 4]. In addition, we examined individual recognition ability based on individual-specific facial features in the three-spined stickleback, along with the significance of red nuptial colouration for male aggressive behaviour. These results of the present study will contribute to resolution of questions regarding the cognitive ability of this model animal.

## 2. Methods

### (a) Animals

The three-spined stickleback *Gasterosteus aculeatus* is widely distributed north of 35°N latitude, where it mainly inhabits coastal areas and lower reaches of rivers [5, 7]. In the breeding season, males establish a breeding territory containing a breeding nest and display nuptial colouration consisting of a bluish dorsal surface and red abdomen [33, 34]. The establishment of a territory precedes nesting [35, 36], and males attack intruders into their territory by adopting a head-down posture and aggressively biting the fins of those intruders [4, 33]. Males become more aggressive to defend their breeding territory during the breeding season than during other seasons, but they have foraging territories in all seasons [37, 38].

For this study, fish were obtained from the sea run population of Shiomi River, Akkeshi, Hokkaido, Japan; bred at Aquatotto Gifu; and kept in aerated stock tanks at 15°C under a 12:12 h light/dark cycle in the laboratory of Osaka City University. The fish were fed commercial flake food (TetraMin; Tetrawerke, Melle, Germany) and bloodworm (Chironomidae, *Propsilocerus akamusi*) once daily. Twenty fish consisting of 10 males (standard length [SL] 60–71 mm) and 10 females (SL 67–79 mm) were used in the experiments conducted from August to November 2020. Due to the constant room temperature and light/dark cycle, the nuptial colours of the males remained present in November although the breeding season had ended in the wild. Fish behaviours were recorded using a video camera (HDR-CX470; Sony, Tokyo, Japan) in all experiments.

### (b) Procedure of Experiment 1

A familiar neighbour was created using an established procedure [26]. Fish were kept in isolation for more than 1 month in stock tanks, and two size-matched fish of the same sex (size difference < 9 mm) were placed in adjacent tanks measuring 18 cm × 30 cm × 24 cm (height) (*n* = 10 males or females in five combinations; each fish had a respective neighbour fish of the same sex) (figure 1). The fish could only make visual contact (figure 2). These experimental tanks contained an air stone and a gardening brick (10 cm × 5 cm × 5 cm) to make the fish more aware of their territoriality.

**Figure 2.**
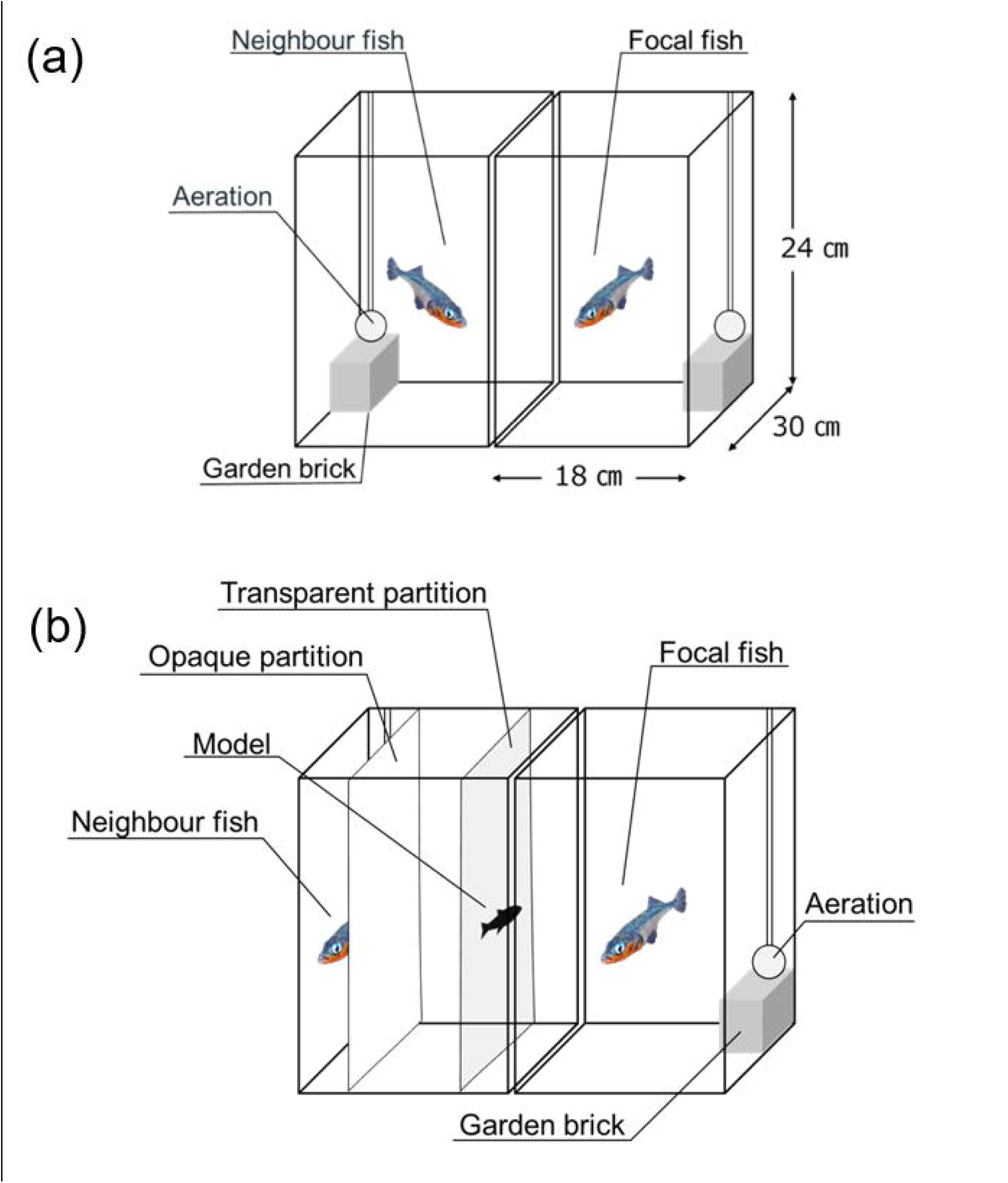
Tank layout for Experiment 1 (*a*) and Experiment 2 (*b*). (a): One fish was introduced in each tank (18 × 30 × 24 cm^3^). Gardening bricks and an air-stone were placed and an opaque partition was removed before the start of experiment. (b): Photographic model on the transparent partition was set after neighbour fish was concealed by an opaque partition. Experiments started after the opaque sheet between two tanks were removed.

In similar previous studies using *Neolamprologus pulcher* [23, 24, 26], the period for establishing the dear enemy relationship between adjacent territorial individuals was set at 5– 7 days. In our preliminary observations, three-spined sticklebacks also showed decreased aggressive behaviour towards adjacent territorial individuals and stabilised their aggression within 7 days, which then increased again when stranger individuals were presented (I. Inoue, personal observation). Therefore, we recorded their behaviours on video for 30 min every 7 days. On day 1, we recorded soon after the partition was removed (09:00), and on days 2 to 7, we recorded around noon. The time for aggressive behaviour of focal fish towards neighbour individuals was analysed for 5 min per day for 7 days. We measured the time for their biting the glass wall facing the neighbouring tank with an open mouth, which are common aggressive behaviours of three-spined sticklebacks against conspecifics [14]. Establishment of a decrease in aggression against familiar neighbours, known as the dear enemy effect [19, 20], is an indicator that animals can discriminate familiar neighbours from strangers or recognise individual familiar neighbours [26, 31]. We placed an opaque partition between the two tanks after lights-out on the last day of Experiment 1.

### (c) Procedure of Experiment 2

Experiment 2 was conducted after Experiment 1, when the neighbours had become less aggressive to each other. In Experiment 2, we acquired photographs of the side view of the 10 familiar neighbours (N) and strangers (S) that had not been encountered by the focal fish for more than 1 month using a digital camera (Nikon D610; Nikon, Tokyo, Japan). Because the red nuptial colouration of males is more distinct when they are face-to-face with other males, we acquired photographs before the end of Experiment 1. These images were edited using GIMP 2.10.28 (The Gimp Team, https://www.gimp.org) to exchange parts of the face between familiar neighbours (N) and strangers (S).

For each focal fish (10 males and 10 females), we created four types of composite photographs with different face/body combinations: face and body of familiar neighbours (neighbour-face and neighbour-body: NfNb), face of familiar neighbour and body of stranger (NfSb), face and body of stranger (SfSb), and face of stranger and body of familiar neighbour (SfNb) (figure 3). We considered the facial area to include the operculum, and we defined the head as indicated in figure 3, similar to the creation of composite photographs in other fish species [22, 24, 26, 28]. Slight adjustment of body colour tone was performed using GIMP to maintain consistent body colouration and to ensure that the margin between the visually transplanted face and body was not obvious to human observers (figure 3). We printed the four models on commercially available glossy photographic paper using an inkjet printer (EP-30VA; Epson, Nagano, Japan); the models were cut to match the contours of the fish and laminated.

**Figure 3.**
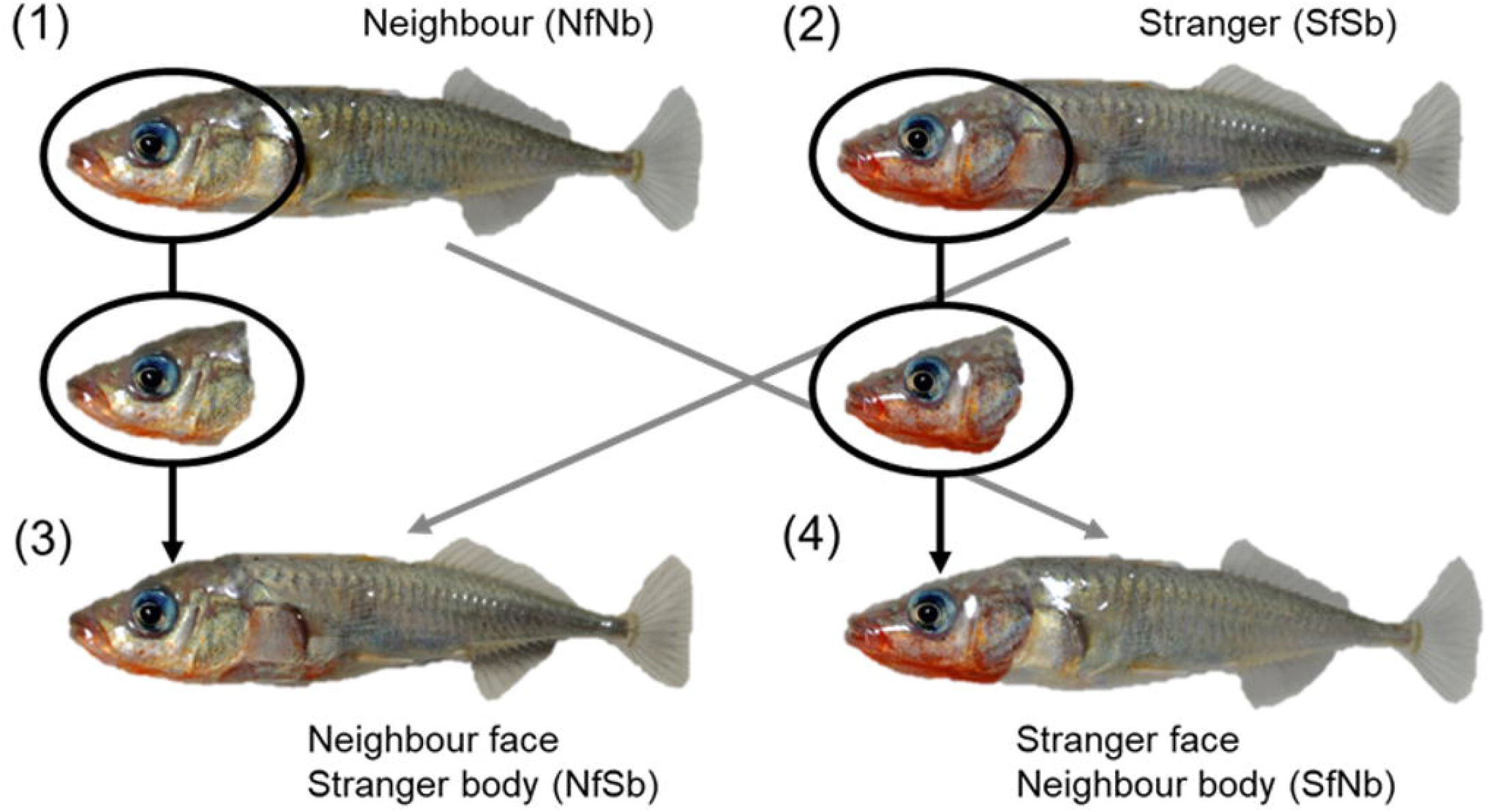
Photographic models presented in Experiment 2. NfNb: the familiar neighbour; SfSb: the stranger; NfSb: face of the familiar neighbour on body of the stranger; SfNb: face of the stranger on body of the familiar neighbour.

We presented one of the four photo-models once every 2 days in random order for 5 min, four times in total over 8 days, at around 12:00 in all cases. During preliminary experiments, focal fish often fled from photo-models attached to the glass wall and remained in the corner of the tank. Therefore, we placed a white opaque partition in the neighbour tank to conceal the neighbour from the focal fish and placed a transparent partition with a photo-model 5 cm from the centre of the boundary of the focal fish tank (figure 2). After 5 min of acclimation when the focal fish had resumed normal swimming, we gently removed the opaque partition between the two tanks and presented the photo-model for 5 min. During the presentation of the model, the behaviour of the focal fish was recorded on video.

We measured the time of aggressive behaviour of the focal fish against the model for 2 min, after the model had been presented and the focal fish had found the photo-model. This time was appropriate because of the gradual decrease in behaviour of the focal fish against the motionless photographs during presentation of the photo-model, consistent with a previous study [26].

Because the focal fish were more aggressive towards strangers than familiar neighbours (see Results), we predicted that the time for aggressive behaviour would be longer in the stranger model (SfSb) than in the familiar neighbour model (NfNb). If three-spined sticklebacks recognise familiar conspecifics only by their facial features, their responses to models of the neighbour-face regardless of body (NfNb, NfSb) would be similar, and responses to models of the stranger-face regardless of body (SfSb, SfNb) would also be similar. In contrast, if these fish use the whole body or features of both face and body to recognise conspecifics, their responses to the modified model (NfSb, SfNb) would presumably be intermediate in comparison to the responses to the unmodified model (SfSb, NfNb).

According to Tinbergen’s studies, aggressive behaviours of male three-spined sticklebacks against photo-models were assumed to be affected by the red area on the belly as a sign-stimulus [4, 10]. The size of the red area considerably varied among subject males (figure 1a,b). Therefore, we measured the area of nuptial red colour between the snout and the base of pelvic fin on the photographs using Adobe Photoshop 23.1.1. (Adobe Systems Inc., San Jose, CA, USA). We then converted the photographs to black and white binary data using ImageJ (http://rsb.info.nih.gov/ij/) and calculated the ratio of red area to the body area excluding the fins. If the key stimulus plays a role in triggering male aggression, focal fish would be expected to show greater aggression towards photo-models with greater nuptial colour.

### (d) Statistical Analysis

Statistical analyses were performed with R version 4.3.2 (R Core Team, 2023) [39]. Linear mixed models (LMMs) in R package *lme4* and *lmerTest* were used for all analyses with individual ID as a random effect to handle the repeated measures design of the within-individual observation. All LMMs were fit by restricted maximum likelihoods, and F-tests with Satterthwaite’s method were applied to test the statistical significance. Effect sizes (partial *η*^*2*^) and their 95% confidence intervals (c.i.) were also calculated using *effectsize* in R package. For post-hoc comparisons, Tukey contrasts were used with the *glht* function in R package *multicomp*. Two-sided *p* values < 0.05 were considered statistically significant for all the tests.

In Experiment 1, we assessed whether the time spent in aggression by focal fish against neighbours of the same sex decreased from day 1 to day 7, by constructing a separate LMM for each sex (*n* = 10 males and 10 females). In Experiment 2, we compared the time spent in aggression by focal fish directed towards four photographic models of the same sex (NfNb, NfSb, SfSb, SfNb) in a separate LMM for each sex (*n* = 10 males and 10 females). Finally, we tested whether the sign stimuli affected aggressive behaviour of focal male against photographic models of the same sex. We constructed a LMM with the time spent in aggression by focal male (*n* = 10) against photograph models as a response variable, photographic model (two models: NfNb, SfSb) as the fixed factor and the ratio of the red area to the total body area as the covariate, including their interaction.

### Data and materials availability

All data are available in the text and Supplementary Materials.

### Ethics

No animals were sacrificed during these experiments. The fish were fed commercial flake food (TetraMin) and bloodworm (*Propsilocerus akamusi*) once daily; they were maintained in a suitable tank environment. Sick or injured fish were removed from the experimental tanks, treated with medication, and only used after recovery. All experiments were conducted in compliance with the Regulations on Animal Experiments of XXX University and the XXX Ethological Society.

## 3. Results

### (a) Experiment 1

The time spent in aggression by focal fish against neighbours of the same sex was highest immediately after the opening of the partition on day 1 (male: 290.48 sec ± 4.56 s.e.m. / 5 min, *n* = 10, female: 227.84 sec ± 15.80 s.e.m. / 5 min, *n* = 10, figure 4). On day 2 and thereafter, the time spent in attack by focal fish decreased significantly in both sexes (male: LMM, *F*_*6,54*_ = 105.69, *p* < 0.0001, partial *η*^*2*^, = 0.92 [95% c.i. = 0.89-1.00], figure 4a; female: LMM, *F*_*6,54*_ = 80.38, *p* < 0.0001, partial *η*^*2*^, = 0.90 [95% c.i. = 0.86-1.00], figure 4b) and stabilized at a lower level (male: <80 sec / 5 min; female: <25 sec / 5 min).

**Figure 4.**
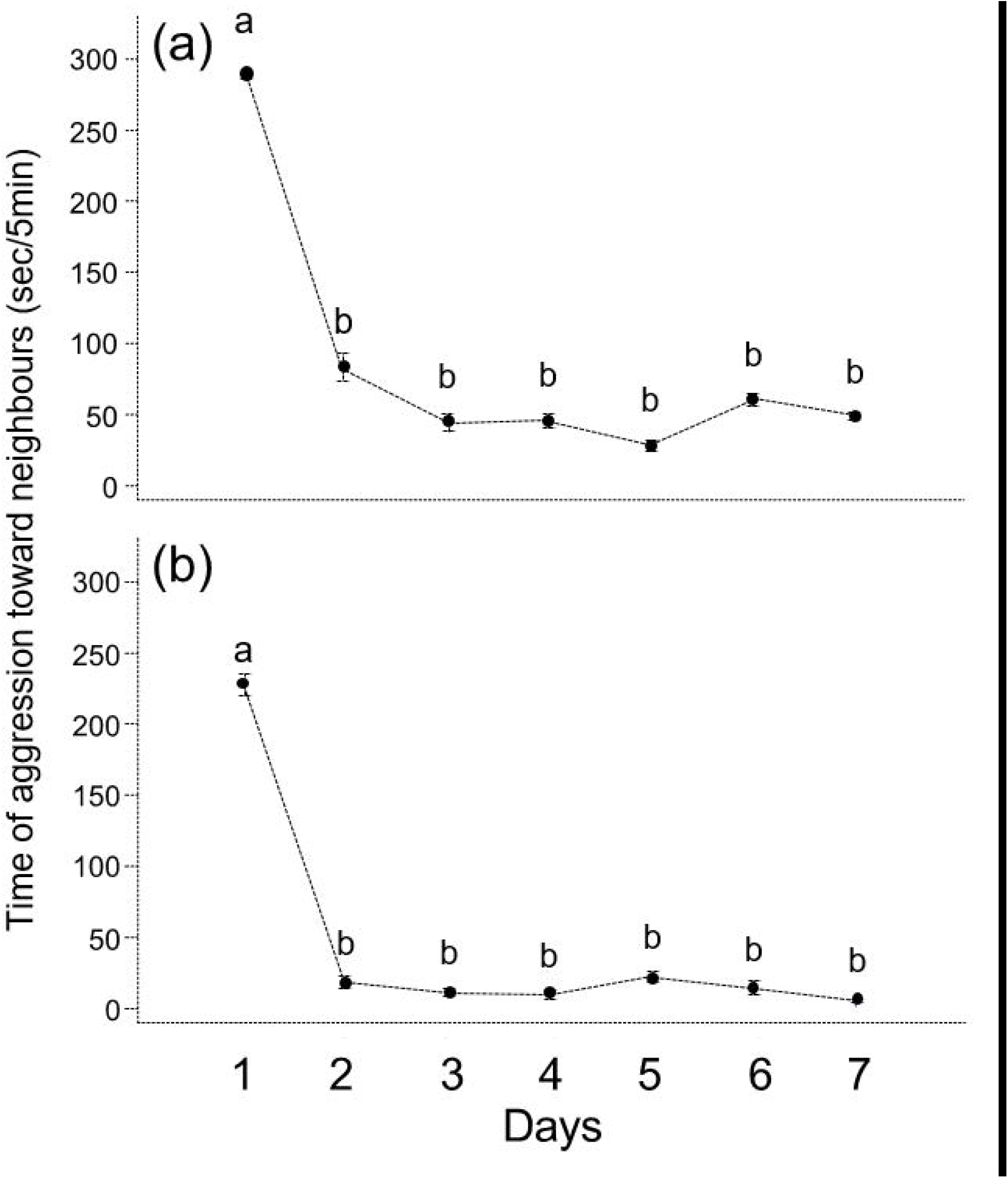
Time spent in attack by focal fish against neighbours of the same sex in Experiment 1. (*a*) Male (*n* = 10). (*b*) Female (*n* = 10). Mean ± s.e.m. a vs b: *p* < 0.05 by Tukey contrasts.

### (b) Experiment 2

Focal fish of both sexes exhibited aggression differently toward four photo-models of the same sex (male: LMM, *F*_*3,36*_ = 12.35, *p* < 0.0001, partial *η*^*2*^, = 0.51 [95% c.i. = 0.29-1.00], figure 5a; female: LMM, *F*_*3,36*_ = 13.59, *p* < 0.0001, partial *η*^*2*^, = 0.53 [95% c.i. = 0.32-1.00], figure 5b). As predicted, focal fish of both sexes spent more time in aggression toward stranger (SfSb) photo-models than toward familiar neighbour photographic models (NfNb). Furthermore, regardless of bodies, focal fish were aggressive significantly and much greater toward both stranger-face photographic models (SfNb and SfSb) than towards both neighbour face photographic models (NfSb and NfNb) (figure 5).

**Figure 5.**
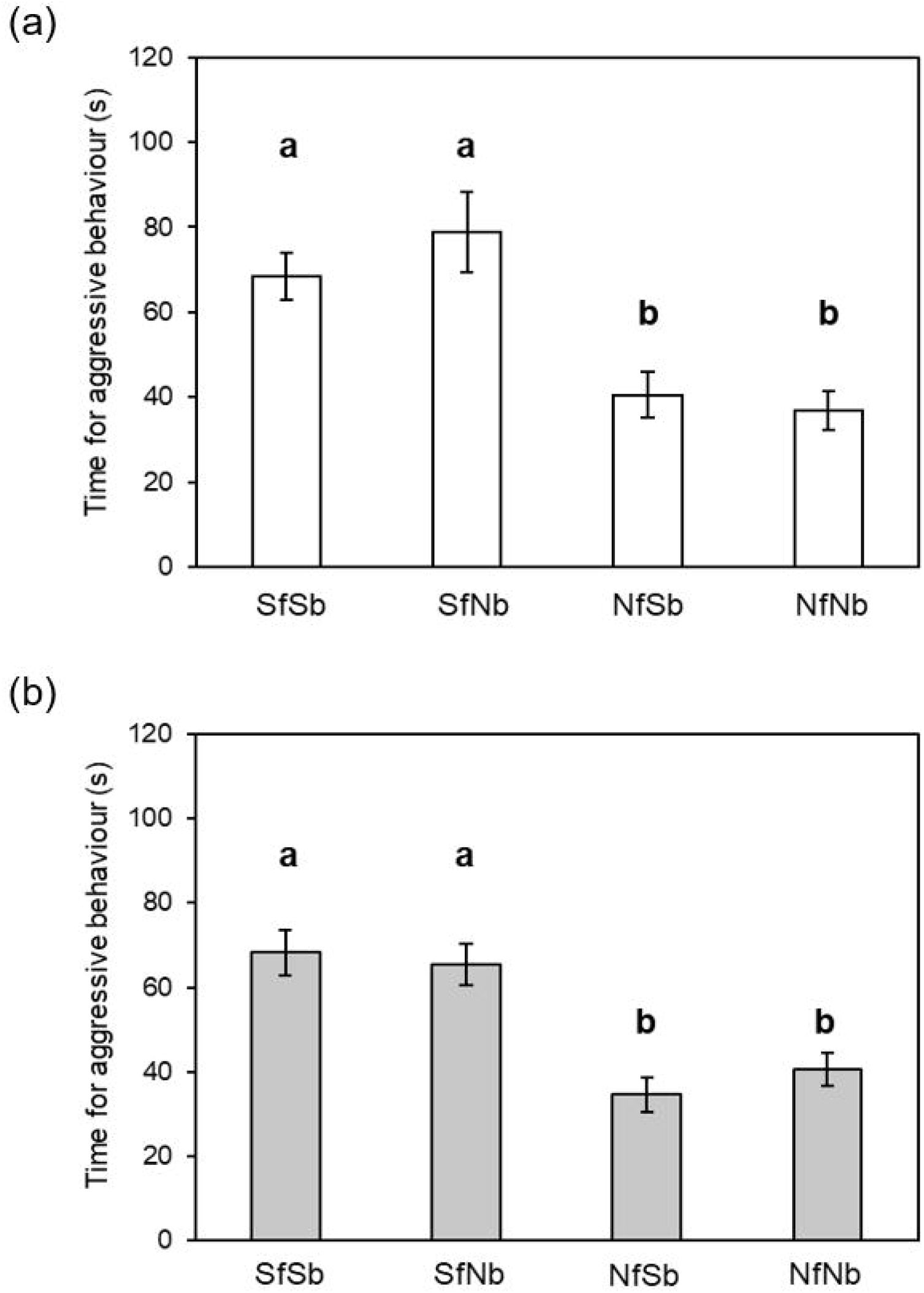
Time spent in aggression by focal fish against four photographic models in Experiment 2. (*a*) Male (*n* = 10). (*b*) Female (*n* = 10). SfSb: the stranger; SfNb: face of the stranger on body of the familiar neighbour; NfSb: face of the familiar neighbour on body of the stranger; and NfNb: the familiar neighbour. Mean ± s.e.m. a vs b: *p* < 0.05 by Tukey contrasts.

Additionally, the ratio of the red area on face to belly to its total body area had no effect on the time spent in aggression by focal males (LMM, photographic model x nuptial colour: *F*_*1,16*_ = 0.22, *p* = 0.65, partial *η*^*2*^, = 0.01 [95% c.i. = 0.00-1.00]; nuptial colour; *F*_*1,16*_ = 0.05, *p* = 0.83, partial *η*^*2*^, = 0.003 [95% c.i. = 0.00-1.00]; photographic model; *F*_*1,16*_ = 12.39, *p* = 0.003, partial *η*^*2*^, = 0.44 [95% c.i. = 0.13-1.00]).

## 4. Discussion

The results of the present study suggested that three-spined sticklebacks of both sexes show dear enemy relationships with their neighbours and considerably reduced aggressiveness towards familiar neighbours. Aggression frequency was much lower against photo-models of familiar faces than against models of stranger faces independent of body type, clearly indicating that three-spined sticklebacks can distinguish familiar neighbours from strangers via facial recognition. Individual recognition based on facial features previously has been reported in fish belonging to the orders Perciformes [23, 25, 27, 29], Beloniformes [30], and Cyprinodontiformes [27]. The three-spined stickleback belongs to the order Gasterosteiformes. These four fish orders are not closely related [40], suggesting that familiar recognition by facial recognition is widespread among teleost fish.

In territorial species, a stranger represents a high level of threat for territory owners, whereas a familiar dear neighbour rarely intruding across the territorial boundary represents a low level of threat (in birds [e.g., 16, 18, 19]; in fish [21–23]). Therefore, current results suggest the three-spined stickleback changes aggression level according to the threat level of adjacent individuals. This suggestion is consistent with previous studies showing that the aggression of sticklebacks is affected by the location of rival males [36]. The rapid identification of individuals around an individual’s territory is indispensable for effective maintenance of a territory by dear enemy relationships [33]. Familiar recognition of dear neighbours has been reported in birds [e.g., 18, 20] and territorial cichlid fishes [23, 25, 26], but these examples solely involved males. Male three-spined sticklebacks have breeding territories and female three-spined sticklebacks have feeding territories [31]; familiar recognition of both sexes will plays an important role in their social lives.

Why do the stickleback and other fishes, including cichlids, guppies, and medaka, distinguish familiar fish based on facial features rather than other body parts? Eyes play an important role in facial recognition among animals and humans (e.g., fish [41, 42], primates [43, 44] and humans, [45, 46, 47]). Studies of eye tracking in primates and humans have shown that they initially tend to focus on an opponent’s face, especially the eyes, before focusing on other body parts (e.g., humans [46], chimpanzees [47, 48], and rhesus monkeys [44]). This suggests that visual cues of individual signals located near the eyes allow rapid signal transmission in these species. Recently, it has been reported that the cichlid *N. pulcher* gazes at the face of encountered fish first and for significantly longer than other body parts [47], consistent with facial recognition patterns in mammals (e.g. [45–47]) and birds [49]. *N. pulcher* can distinguish between familiar and unfamiliar faces within 0.5 s with high accuracy [23]. For this type of rapid and accurate facial recognition, individual-specific signals near the eye (i.e., individual variation in face colour morphs (figure 1a, b)) should also be effective in the three-spined stickleback.

Three-spined sticklebacks can discriminate a familiar neighbour from a stranger based on individual faces. The facial feature represents an individual-specific signal, and this facial recognition process is similar to the process of true individual recognition (TIR) rather than class-level recognition [25, 31, 50]. The cichlid *N. pulcher* that can distinguish familiar dear neighbours from stranger fish was able to identify two familiar dear neighbours by their specific facial features (i.e., TIR) [23, 26]. We predict that the three-spined stickleback, a species with individual-specific faces, can also perform TIR [32]. The ability to recognise multiple dear neighbour fish (i.e., TIR) is necessary to effectively maintain stable territoriality. This cognitive ability is likely to be highly sophisticated, probably similar to mammals [32]. Our observations suggest that sticklebacks that can recognise and respond appropriately to a familiar individual will have complex cognitive capacity, which could be associated with mental mechanisms different from the innate releasing mechanism with sign-stimulus.

The innate releasing mechanism predicts that territorial aggression of stickleback males will be affected by the size and brightness of the red nuptial colour (i.e., the sign-stimulus) [3, 4]. In the present study, however, we observed cases in which four males lacking red colour received intensive aggression on day 1 in Experiment 1; the size of the red area, including its absence, had no effect on the intensity of male aggression in Experiment 2 (figure 1b). This was consistent with previous reports that the degree of red nuptial colouration on the belly of this fish had no effect on the intensity of territorial aggression [10, 11]. However, it should be noted that since the models presented here were photographs, it is not possible to comment on the effect of chemical stimuli and the probability of winning or intimidating effect that the red belly might influence [12]. Furthermore, females lacking red colour exhibited intensive aggression against non-familiar females. In any case, our observations do not support the interpretation of aggression prompted by the “sign-stimulus”.

Milinski [13] reported that three-spined sticklebacks have a tit-for-tat strategy during repeated predator inspection visits. Such a strategy requires TIR of the opponents and a strong memory [51]. The results of the present study provided evidence for the ability of familiar recognition and suggest TIR in sticklebacks. A recent study showed no evidence of familiar recognition in the three-spined stickleback [15]; however, lack of evidence of a capacity does not always constitute evidence of its absence. Perhaps, considering the present results and the ecology of sticklebacks, they are able to associate rocks and food, but no relationship could be found between the recognised individuals and food. Because the former situation is their daily life, but the latter situation is rare. Many examples of high cognitive abilities in social fishes have been reported thus far, including prediction of the behaviour of others based on an individual’s own experience during coordinated hunting [52], transitive inference in social dominance [53, 54], use of prosocial and antisocial choices [55], and mirror self-recognition [56-58]. These sophisticated forms of social cognition and maintenance of stable sociality, such as dominance order and territoriality, require the ability to recognise specific individuals. The present findings suggest that the three-spined stickleback represents the evolution of some intelligence or cognitive abilities that is not explained as responds against sign stimuli in fishes within the context of complex social relationships [59].

## Notes

### Competing Interest Statement

The authors have declared no competing interest.

